# Leveraging protein language models and scoring function for Indel characterisation and transfer learning

**DOI:** 10.1101/2025.03.12.642715

**Authors:** Oriol Gracia I Carmona, Vilde Leipart, Gro V Amdam, Christine Orengo, Franca Fraternali

**Affiliations:** Department of Structural and Molecular Biology, University College London, London, UK; Randall Centre for Cell & Molecular Biophysics, King’s College London, New Hunt’s House, Guy’s Campus, London, UK; Faculty of Environmental Sciences and Natural Resource Management, Norwegian University of Life Sciences, Ås, Norway; School of Life Sciences, Arizona State University, Tempe, Arizona, USA

**Keywords:** Transfer learning, zero-shot inference, protein language models, interpretability, indels, pathogenicity predictors

## Abstract

1.

Protein language models (PLMs) are increasingly used to assess the impact of genetic variation on proteins. By leveraging sequence information alone, PLMs achieve high performance and accuracy and can outperform traditional pathogenicity predictors specifically designed to identify harmful variants contributing to diseases. PLMs can perform zero-shot inference, making predictions without task-specific fine-tuning, offering a simpler and less overfitting-prone alternative to complex methods. However, studying in-frame insertions and deletions (indels) with PLMs remains challenging. Indels alter protein length, making direct comparisons between wildtype and mutant sequences not straightforward. Additionally, indel pathogenicity is less studied than other genetic variants, such as single nucleotide variants, resulting in a lack of annotated datasets. Despite these challenges, approaches that leverage PLMs through transfer learning have emerged, making it possible to capture the features needed for more accurate predictions. Still, the current approaches are limited in terms of allowed organisms, indel length, and interpretability. In this work, we devise a new scoring approach for indel pathogenicity prediction (IndeLLM) that provides a solution for the difference in protein lengths. Our method only uses sequence information and zero-shot inference with a fraction of computing time while achieving performances similar to other indel pathogenicity predictors. We used our approach to construct a simple transfer learning approach for a Siamese network, which outperformed all tested indel pathogenicity prediction methods (Matthews correlation coefficient = 0.77). IndeLLM is universally applicable across species since PLMs are trained on diverse protein sequences. To enhance accessibility, we designed a plug-and-play Google Colab notebook that allows easy use of IndeLLM and visualisation of the impact of indels on protein sequence and structure. The tool is available on GitHub https://github.com/OriolGraCar/IndeLLM and Colab https://colab.research.google.com/drive/1CgwprttaNFR_KeJGyFzP0a0C9Y wc4P.

**Graphical Abstract, if needed, or logo til include on Google Colab:** 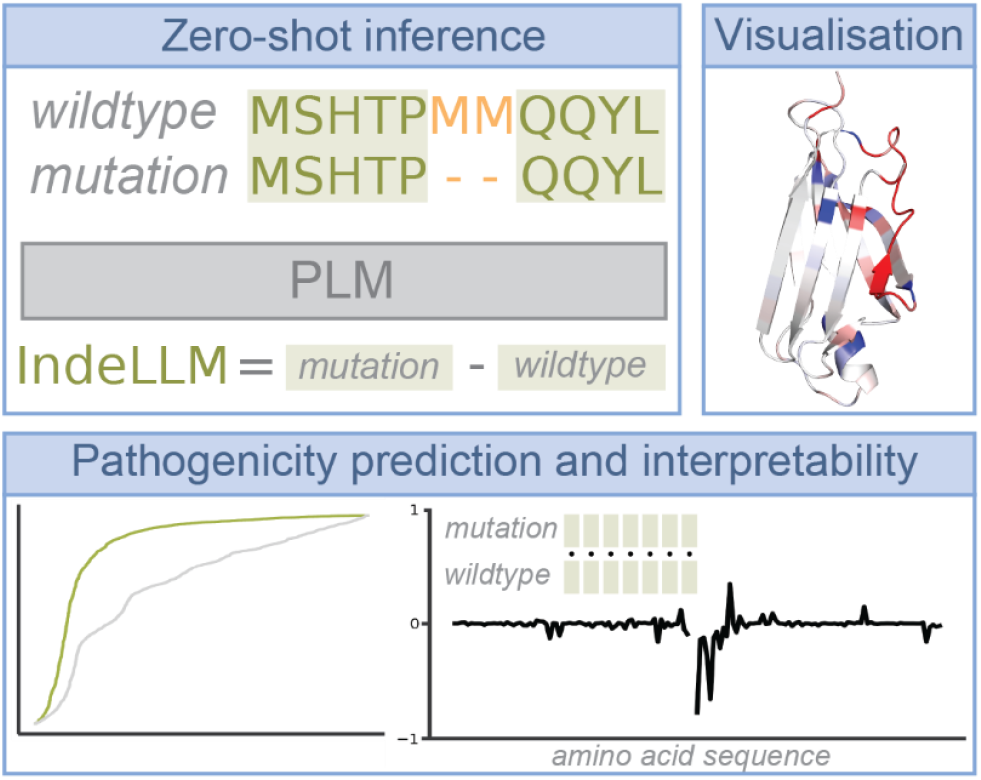

## 2. Introduction

Insertion and deletion variants (indels) are among the most prevalent forms of genetic variation, accounting for about 18% of all variation in humans, with individuals sometimes carrying hundreds of these mutations across their exomes [1,2]. These variations are commonly categorised into in-frame indels, which add or remove entire codons without disturbing the reading frame, and frameshift indels, which shift the reading frame by adding or removing nucleotides in non-triplet amounts, producing entirely new downstream protein sequences. The effects of indels can vary significantly, with some leading to substantial changes in protein structure and function while others remain largely benign [3,4].

Frameshift indels, which disrupt the reading frame and typically result in loss of protein function, are generally considered pathogenic [5]. Here, we are considering in-frame indels, which, in contrast, preserve the reading frame while altering the amino acid sequence. These indels can potentially impact protein stability and function. Their effects are less predictable than frameshift indels and are often comparable to missense variants [6].

Although in-frame indels are known to cause several well-documented monogenic diseases, such as cystic fibrosis, eye disorders such as childhood cataracts and retinal dystrophies, and several types of human cancers [5–7] their pathogenicity has historically been less studied than that of Single Nucleotide Variants (SNVs) [1,8].

As a result, the mechanism behind pathogencity for in-frame indels is far from understood. Despite this, some specialised computational approaches are available to predict in-frame indel pathogenicity. The tools employ diverse strategies, often relying on the use of several manually encoded predictive features, such as sequence conservation, indel size and protein structure and function, that have been seen to correlate with pathogenicity [8–12]. However, selecting such features is based on an incomplete knowledge of the underlying mechanisms. It may not accurately capture the full complexity of indel pathogenicity or potentially introduce biases that emphasise certain aspects while overlooking others. In addition, most available tools are often tailored to the human genome, as creating more broadly applicable tools is challenging due to limited available pathogenicity annotations for indels from other species.

Protein Language Models (PLM) have the ability to meet these challenges. These models have emerged as a transformative approach in artificial intelligence, particularly for specialised fields like protein sequence analysis. PLMs are able to capture intricate relationships in proteins only by using protein sequences as input through training on massive unlabeled protein sequence datasets. The obtained embeddings contain rich biological information which can be used as input for transfer learning approaches.

These PLMs can capture complex information about protein secondary and tertiary structures across a wide range of organisms, from bacteria to humans, using the amino acids sequence as its only input feature, potentially providing insights beyond those available from traditional methods at a faster speed. Advances in computational power and the growth of available protein sequences drive the field of PLMs to create models with advanced transformer architectures containing billions of parameters. Today, a diverse range of PLMs, varying in sizes and architectures, is available, from general-purpose to specialised models [13–18].

Transformer-based architectures and PLMs have shown great success and revolutionised the SNVs prediction field [13,19,20], both when used as zero-shot inference methods (using the predicted logits with no additional training [21]) and when fine-tuned or used as features for other models. Zero-shot inference approaches are appealing since they do not risk overfitting to the data. However, when dealing with indels, using these techniques is not straightforward because of the difference in length between the sequences, which adds extra challenges [19,22]. Recent efforts have focused on developing new methods leveraging PLMs, either as zero-shot inference tools using a generalised scoring function described by Brandes et al. or as features to train other downstream models [9,23]. However, these approaches still have limits in their accuracy or possible use cases. For example, SHINE, an elastic net model trained on a principal component decomposition of PLMs embeddings, is limited to small in-frame indels of one to two amino acids and requires two independent models, one for deletions and one for insertions [23].

Despite these efforts, it remains unclear which PLMs provide the most accurate latent representations of the protein effects caused by indels, how best to transfer those representations, which scoring functions perform best for zero-shot indel prediction, and the limitations of these methods. Furthermore, the interpretability of predictions from PLMs can be challenging, as the patterns they capture are often complex and not immediately intuitive [24,25].

In this work, we present an in-depth analysis of the inner workings of general scoring functions for zero-shot pathogenicity classification for in-frame indels. We suggest an improved scoring regime that achieved an improvement of 0.26 points in the Matthews correlation coefficient (MCC) over a general scoring approach [19], reaching an MCC comparable with the best-performing methods for indel prediction without any further training. From there, we test different transfer learning approaches on a simple one-hidden-layer Siamese network and manage to obtain unprecedented performance through a biology-guided slicing of the PLM embeddings. Finally, we present an analysis approach that aids in understanding the model predictions by looking at the local effects of the indel on the rest of the protein amino acids. The approach, named IndeLLM, is easy to use through a Google Colab version https://colab.research.google.com/drive/1CgwprttaNFR_KeJGyFzP0a0C9Y wc4P, and the code is freely accessible on https://github.com/OriolGraCar/IndeLLM.

## 3. Methods

### Dataset Generation (including handling of long sequences)

We retrieved datasets of indels in the human genome from three recent studies on indel predictions [9,19,23]. The first study, Cannon et al., evaluated the performance of nine in silico pathogenicity prediction tools for small in-frame indels [9]. Their published dataset contains 3964 human indels (genome build GRCh38) collected from gnomAD, ClinVar, and the Deciphering Developmental Disorders (DDD) study [26–28] with predictions classifications from nine tools: CADD, CAPICE, FATHMM-indel, MutPred-Indel, MutationTaster2021, PROVEAN, SIFT-indel, VEST-indel and VVP [8,10,11,29–34]. The Fan et al. study presented a protein language model-based (ESM1b) pathogenicity predictor for short in-frame indels called SHINE [23]. We retrieved the 1-2 amino acid indel training and validation dataset for SHINE. The Fan et al. dataset contains 5457 indels from ClinVar and gnomAD. The Brandes et al. study included a dataset that was used to benchmark the performance of a PLM (ESM1b) on in-frame indels predictions, which is part of a genome-wide disease variant effect prediction study [19]. The Brandes et al. dataset contained 3470 ClinVar indels.

The Cannon et al. dataset did not contain the variant impacts on protein level, so we submitted the reported chromosome position and nucleotide variation to Ensembl Variant Effect Predictor (VEP [35]) to obtain the reference and alternative amino acids for most indels (some transcription IDs were no longer available). Then, we used the available transcription IDs (cross-referenced with the output from VEP) to obtain the full-length peptide sequence from Ensembl BioMart [36]. We quality-checked that the dataset only contained in-frame indels, leading to a single protein impact per indel, excluding frameshift variants. The Fan et al. dataset contained Protein IDs, which we submitted to BioMart to obtain the full-length peptide sequences (some protein IDs were no longer available). The Brandes et al. dataset included the cropped wildtype and mutated protein sequences that we could use directly in our study.

We combined all the unique indels from the three studies creating our dataset (n = 7500) available at https://github.com/OriolGraCar/IndeLLM. The dataset includes 7500 indels, where 2409 are insertions and 5091 are deletions. Of these, 2878 are classified as likely pathogenic or pathogenic, while 4622 are classified as likely benign or benign. The indels range from 1 to 223 in amino acids inserted or deleted (length) and represent a distribution of short indels (1-2 amino acids, n = 3882) and longer indels (>= 3 amino acids, n = 3618) (Supplementary Material section 1 for more details). We used this dataset for all analyses, except the specific performance analysis comparing our scoring approach to the nine indel prediction tools, in which we use the Cannon et al. subset of the dataset. Cannon et al. included the reported scores from the nine prediction tools. The Cannon et al. subset has 3478 indels from gnomAD (n = 647), ClinVar (n = 2577), and DDD (n = 254), where 1115 are insertions and 2363 are deletions. Of these, 1518 are classified as likely pathogenic or pathogenic, while 1960 are classified as likely benign or benign. The subset is labelled in our published extended dataset, available at https://github.com/OriolGraCar/IndeLLM. Based on this specific performance analysis, we found that Provean [32] and MutPred-indel [11] showed no signs of overfitting and are among the few methods that have not been discontinued. We collected prediction scores for all the 7500 indels using Provean and MutPred-Indel, which are included in our dataset.

Due to the size limitations of some PLMs, it is not possible to fit sequences with more amino acids than the allowed number of tokens. Models that allow for sequences of any length, such as ESM2, which uses RoPe positional encoding, quickly run into memory limitations due to the memory required to grow quadratically with protein length. Because of these, sequences exceeding 1000 amino acids were truncated around the indel location, leaving a buffer region of 500 amino acids down and upstream of the indel. Since the attention mechanism grows quadratically with sequence length, most available PLMs have limits on the distance between tokens for which attention is computed, usually capped at 500 amino acids [1,2]. By cropping the sequences and leaving 500 amino acids around the indel location, we guarantee that the maximum amount of the learned self-attention can be used for the effect inference. The resulting cropped sequences have a limit size of 1000 amino acids plus the length of the indel. In cases where the indel length exceeds the models’ size limit, the number of buffer amino acids around the indel is symmetrically reduced to a length that fits the model (1022 amino acids). All used sequences in full length are provided in the dataset, while their respective crops and annotations are provided at https://github.com/OriolGraCar/IndeLLM.

### Indel scoring

Brandes et al. described a generalised approach for scoring in-frame indels as a zero-shot inference task [19]. In summary, the described approach calculates a pseudo-log likelihood (PLL) of each sequence s as:

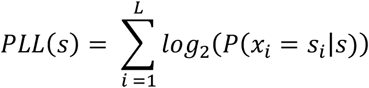

Where L is the sequence length, s_i_ is the amino acid at position i, and *log*_*2*_*(P(x*_*i*_ *= s*_*i*_|*s))* is the log-likelihood predicted by ESM1b for observing the input amino acid s_i_ at position i given the entire input sequence s. In this context, the output of the PLMs is treated as a sequence of random variables x = x_1_,…,x_l_, where each x_i_ represents the probability distribution for observing one of the 20 standard amino acids at position i. The effect score of an in-frame indel is calculated as the difference in pseudo-log-likelihoods between the mutated (s_mut_) and wildtype (s_WT_) sequences: PLL(s_mut_) − PLL(s_WT_). Since the scoring above calculates the pseudo-log-likelihood of the whole sequence, which was first presented in the Brandes et al. study [19], we refer to this scoring as “Brandes.” One limitation of such a scoring method is that it relies on subtracting incomparable values due to the length difference of the sequences, especially when dealing with longer indels. The study showed attempts to limit that effect using a scaling factor dependent on the length difference but showed no improvements compared to the original Brandes score [19]. In this work, we present an alternative scoring approach to obtain comparable PLLs by calculating the PLL of only the overlapping amino acids. Since PLMs are context-aware language models, the indels’ effect is observable on the marginal probabilities of the rest of the amino acids in the sequence, allowing for the subtraction of comparable PLLs. The scoring can be computed as before, and the PLL of the overlapping regions (PLL_overlap_) can be calculated as:

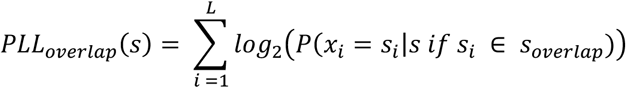

Where s_overlap_ is the amino acids present in both the wildtype and mutated sequences, we refer to this scoring as IndeLLM. The optimal pathogenicity threshold for the entire dataset using IndeLLM is −0.59. This threshold was determined by maximising the difference between the true and false positive rates on the receiver operating characteristic (ROC) curves.

Finally, we also compare the predictions’ performance when using a masking objective for the assessed position or when using inference without masked tokens.

### Siamese network architecture

For the Siamese network, we split the dataset into training, validation, and test datasets containing 80%, 10%, and 10% of the indels, respectively. We used CD-HIT [37] to cluster wildtype sequences with a maximum 50% sequence identity (2902 clusters), and the sequences in each cluster were always kept in the same dataset so that our training, validation, and test datasets contained diverse sequences. The clusters were randomly distributed into the three datasets while ensuring close to equal fractions of indel types, lengths, and pathogenicity classification as in the entire dataset. We considered indels of 1-2 amino acid short, while the remaining long. See Table 1 for the distributions per dataset.

**Table 1:** The distribution of short and long deletions and insertions and the number of benign and pathogenic indels for the full dataset and the splits (training, validation and test).

| Dataset | Total indels | Deletions |  | Insertions |  | Benign | Pathogenic |
| --- | --- | --- | --- | --- | --- | --- | --- |
|  |  | Short | Long | Short | Long |  |  |
| **Full** | 7500 | 3374 | 1717 | 1534 | 875 | 4622 | 2878 |
| **Training** | 5960 | 2690 | 1357 | 1225 | 688 | 3774 | 2186 |
| **Validation** | 819 | 367 | 194 | 161 | 97 | 453 | 366 |
| **Test** | 721 | 317 | 166 | 148 | 90 | 395 | 326 |

To enhance indel classification and explore transfer learning potential using PLMs, we implemented a shallow feed-forward neural network (FNN) inspired by the architecture described in VariPred for SNVs [20]. The FNN was trained solely on the class labels, keeping the PLM parameters frozen. A simple grid search was employed for hyperparameter optimisation, with the final parameters for each model detailed in Figure 2. To mitigate overfitting, the FNN was designed with a single fully connected hidden layer. Each model was trained five times with different initial weights to evaluate reliability.

The input features extracted for the models were as follows: mean embeddings of the last hidden layer for the wildtype sequence, mean embeddings of the last hidden layer for the mutant sequence, mean embeddings of the last hidden layer for the deleted or inserted amino acids, the IndeLLM score, the length of the indel, and the type of indel (0 for deletion, 1 for insertion). For models utilising indel embeddings, the embeddings corresponding to the indel region were excluded from the longest sequence’s embeddings to ensure that only corresponding regions of the protein were compared.

Four final model architectures were developed, each leveraging a different combination of these features:

- **Model 1:** A basic Siamese network that used the mean embeddings from the last hidden layer of the PLM for the whole wildtype and mutant sequences.
- **Model 2:** Built on Model 1 by adding the IndeLLM score as an additional input parameter.
- **Model 3:** Extended Model 2 by incorporating the indel type (insertion or deletion) and indel length as additional parameters.
- **Model 4:** This model took a more advanced approach by splitting embeddings into overlapping and non-overlapping regions. The inputs included the mean embeddings from the overlapping regions of the wildtype and mutant sequences, the mean embeddings of the indel (non-overlapping region), the indel type, and its length. This biologically informed architecture aimed to capture sequence structure better.

Each model was evaluated to determine the contribution of its specific features to the indel classification task. The sequence embeddings were extracted using ESM2 with 650M parameters. A final output layer with two nodes and a SoftMax activation function was used to classify variants as pathogenic or benign. Only the pathogenic node’s output was considered for classification. The area under the curve (AUC) on the validation dataset was used as the early stopping criterion during training. The best-performing pathogenicity threshold for all the data is 0.46.

### Performance analysis

To compare the performance of the pathogenicity prediction tools and PLMs, we computed the ROC curve, F1 score, and Matthews correlation coefficient (MCC) score and confusion matrixes using Scikit-learn [38]. The ROC curve provided the false and true positive rates (FPR and TPR) at different threshold values. When comparing any approach in our study, we used the calculated FPR and TPR to find the optimal threshold, rather than the default threshold provided by each pathogenicity prediction tool, to make a fair comparison. The FRP and TPR were also used to calculate the ROC curve’s AUC. Our calculations are provided in our analysis Jupyter Notebooks at https://github.com/OriolGraCar/IndeLLM.

### Interpretability analysis

Since PLMs are context-aware, one can compare the changes of the predicted amino acid probabilities at each position due to the surrounding context by subtracting the individual probabilities from each other for the amino acids that are overlapping:

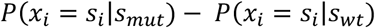

Where (S_i_|S_mut_)is the first amino acid in the mutated sequence S_mut_, and (S_i_|S_wt_) is the same amino acid but in the wildtype sequence S_wt_. This results in a different score for each position in the sequence.

These localised changes in probabilities can then be mapped back to the 3D structure of the protein of interest. This allows one to observe the regions where the model predicts the bigger changes due to the indel and understand which positions drive the pathogenicity prediction results.

To illustrate this approach, we identified genes in our dataset with multiple indels reported, both pathogenic and benign. We selected two proteins, Fibroblast growth factors receptor 1 (FGFR1) and Glomulin (GLMN), which had their 3D protein structure resolved at high resolution. We show two domains of FGFR1, the apo structure of the intracellular tyrosine kinase domain (PDB ID 4UWY [39] Fig 5E) and the first of the three extracellular Ig-like domains (PDB ID: 2CR3, Fig 5F), and the complete GLMN chain extracted from the Glomulin-RBX1-CUL1 Complex (PDB ID: 4F52 [40], Fig 5I). The probability changes (difference scores) are obtained by aligning the wildtype and mutated sequence using Bio.AlignIO package is from Biopython [41] and then coloured on the structure using PyMOL [42].

## 4. Results

### 4.1 Efficient scoring approach for zero-shot inference using Protein Language Models

Our dataset has 7500 indels, where 2409 are insertions, and 5091 are deletions. Of these, 2878 are classified as likely pathogenic or pathogenic, while 4622 are classified as likely benign or benign. We used the dataset to asses different scoring methods for the zero-shot inference using PLMs. The PLM of choice to test the scoring methods was ESM2 650M parameters [15], which have shown the best performance compared to the other tested PLMs (Supplementary Material section 2). Using PLMs as zero-shot predictors is not straightforward due to the varying lengths of both the sequences and indel sizes [22]. Brandes et al. [19] described a generalisable approach to score sequences using the pseudo-log-likelihood (PLL) of each sequence calculated as the sum of the probabilities of each observed amino acid (see methods for more details). The described general approach, referred to as Brandes scoring, achieved a reasonable performance of MCC 0.39 (Fig 1A-1B).

**Figure 1:**
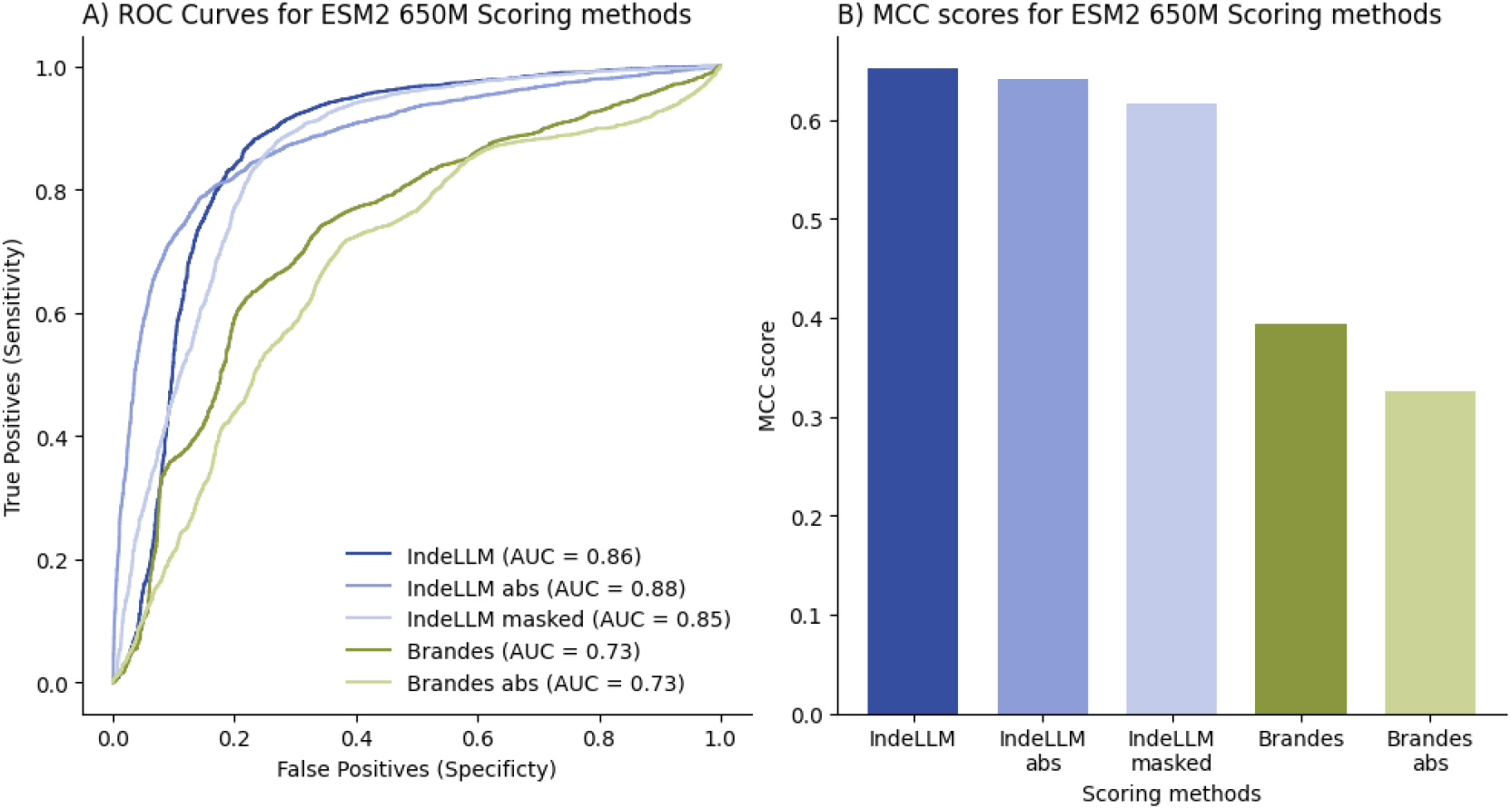
Evaluation of different scoring methods using the probabilities of ESM2 (650M parameters). The different scoring methods were IndeLLM (probabilities of overlapping regions), IndeLLM abs (using the absolute probability of overlapping regions), IndeLLM masked (logits from masked positions), Brandes (sum of all probabilities) and Brandes abs (using the absolute sum of probabilities). We compared the scoring methods by A) plotting the ROC curve and calculating the AUC and B) calculating the MCC scores. See Supplement File 1, Table S1 for all scores.

**Figure 2:**
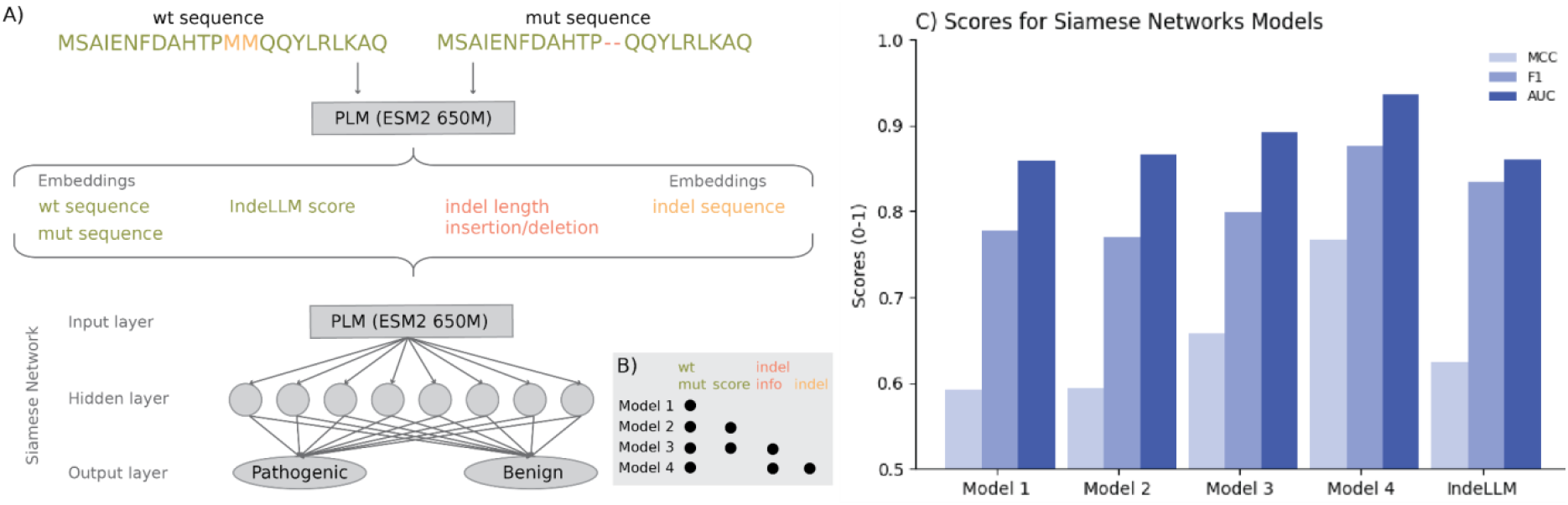
A) Siamese network setup. From the wildtype and mutated sequence, we generated 4 types of weights: The mean embeddings for the wildtype and mutated sequence (the full sequences), the IndeLLM score (using only overlapping regions, coloured green), the indel type and length (based on the gap position and size, coloured red) and the mean embeddings of the inserted or deleted amino acids (indel sequence, coloured orange). The setup includes only one hidden layer of 8 nodes. B) Table of the weight used in Model 1-4 marked with black dots and the same colouring as in panel A. C) The MCC (light blue), F1 (blue) and AUC (dark blue) scores for the mean of Model 1-4 and IndeLLM using the test dataset.

One concern of such a scoring method is that the PLLs being compared are not comparable due to the difference in length in the wildtype and mutated sequences. To mitigate this, we removed the probabilities of the deleted or inserted amino acids, calculating a PLL for the overlapping regions. Doing this eliminates the issue of incomparable likelihoods by indirectly inferring the indel’s effect. This scoring approach, which we denoted as IndeLLM, produced an improvement in the predictions, ranking first among the tested scoring methods (Fig 1A-1B and Supplement File 1, Table S1).

Another factor to consider is the impact of indels, which significantly increase the likelihood of a protein being potentially pathogenic. To account for that, the absolute value of the difference between PLL is used instead. However, this scoring method, denoted as IndeLLM abs or Brandes abs, showed no improvement compared to just using the plain subtraction of PLLs (Fig 1A-1B).

Finally, we also compared two different ways to calculate the marginal probabilities, one using a single encoding step and then using that to compute the logits at each position (IndeLLM). The other uses a masking objective by masking each position individually and then using the logits of the masked position only (IndeLLM masked). Both approaches showed indistinguishable performances (Fig 1A-1B, MCC IndeLLM: 0.65 vs. IndeLLM masked: 0.62), with the single encoding approach requiring significantly less computation time.

### 4.2 Leveraging transfer learning and Siamese networks for Indel prediction

We leveraged PLMs in combination with a Siamese network to predict the pathogenicity of Indels. Since PLMs are pre-trained models, we can extract high-quality embeddings that encode the structural and functional properties of sequences without requiring extensive task-specific training. These embeddings were fed into smaller, task-specific architectures, a Siamese network, to efficiently compare and analyse sequence pairs.

Due to the small size of the labelled indels dataset (n = 7500), we used a small multilayer perceptron (MLP) model with only one hidden layer. This mitigated overfitting by limiting the model’s complexity and, thus, its capacity to memorise the training data by picking up patterns in the noise that are not generalisable to the populations. In particular, using small Siamese networks for pathogenicity prediction has shown high performance in single-point mutation pathogenicity prediction [20].

We used 80% (n = 5960) of the labelled indels for the training dataset, while the remaining indels were split into a validation and test dataset (n = 819 and n = 721, respectively). A robust model should be able to predict the pathogenicity of unseen protein sequences and indels. Therefore, it is essential to avoid redundancy between training, test and validation datasets (data leakage). We clustered all wildtype protein sequences by sequence identity above 50% and randomly divided the clusters into the three datasets, achieving a maximum of 50% identity between all three datasets. We also ensured that the distribution of indel type (insertions and deletions), indel size (short (1-2 amino acid) or long indels (>= 3 amino acids), and classification (pathogenic and benign) were equal to the distribution in the complete dataset (see method for more details).

We used a similar setup to the one suggested by VariPred [20]. Namely, a Siamese network with one hidden layer, using a LeakyReLu as an activation function and a dropout rate of 0.5. Other hyperparameters, such as the number of neurons in the hidden layer, batch size, learning rate and the negative slope parameter for the LeakyReLu, were optimised through a simple grid search (Fig. 2A).

The difference in length between the wildtype and mutated sequences adds an extra layer of complexity when deciding which part of the embedding to use as input. Other approaches using PLM embeddings as input features have solved this by separating insertions from deletions [23]. Here, we tested 4 different Siamese architectures to find which approaches provide the best transfer of the relevant latent space (Fig 2A-2B).

- Model 1: Siamese network with mean embeddings of the last hidden layer for wildtype and mutant sequences.
- Model 2: Same as Model 1, but with the addition of the IndeLLM score as an input.
- Model 3: Same as Model 2, but adds indel type (insertion or deletion) and length as extra inputs.
- Model 4: Siamese network where embeddings are split, using mean embeddings of overlapping and non-overlapping (indel) regions, with indel type and length as extra inputs.

All replicates for Model 1 using a more classical representation of the wildtype and mutant sequence by extracting the mean embeddings performed equally to the IndeLLM score in terms of AUC of the ROC curve but generally underperformed in terms of MCC and F1 score (Performance on the test dataset was AUC: 0.86 vs 0.86, MCC: 0.59 vs 0.62 and F1: 0.78 vs 0.83, respectively), indicating that the mean embeddings did not contain unique information when compared to the proposed IndeLLM score. Model 3 was the only model from the group with a comparable MCC to the IndeLLM score (0.66 vs. 0.62, respectively), suggesting that providing the indel length and type contributed information on top of the plain mean embeddings. On the other hand, Model 4 performed significantly better than the IndeLLM score and any of the other 3 models, achieving MCC performances on the test set of 0.77, a 24% improvement from the IndeLLM score. This suggests that the information captured by the embeddings of the inserted or deleted amino acids adds additional information to the sequence embeddings. A more detailed breakdown of the performances of each model on each of the available datasets can be found in Supplementary File 1, Table S2.

### 4.3 Comparing Indel predictor performance to the IndeLLM scoring approach and siamese model

We used the scores of nine different tools published by Cannon et al. [9] and evaluated their performance and indications of overfitting (Supplement Material sections 3 and 4). We concluded that the best-performing indel pathogenicity prediction tools were Provean and MutPred-indel, which were used to obtain pathogenicity prediction for all 7500 indels. We compared performance using the IndeLLM score to the Provean and MutPred-indel on all indels (n=7500), the training (n=5960), validation (n=819) and test (n=721) datasets. We observed consistent pathogenicity prediction performance for all methods (Fig 3). MutPred-Indel and IndeLLM have comparable performance (MCC MutPred-Indel: 0.57 vs MCC IndeLLM: 0.62). IndeLLM Siamese outperformed all methods and has 5.5% better prediction accuracy than the best-performing prediction tool Provean (MCC IndeLLM siamese: 0.77 vs MCC Provean: 0.73) (Fig 3F). See Supplementary File 1, Table S3 for all scores.

**Figure 3:**
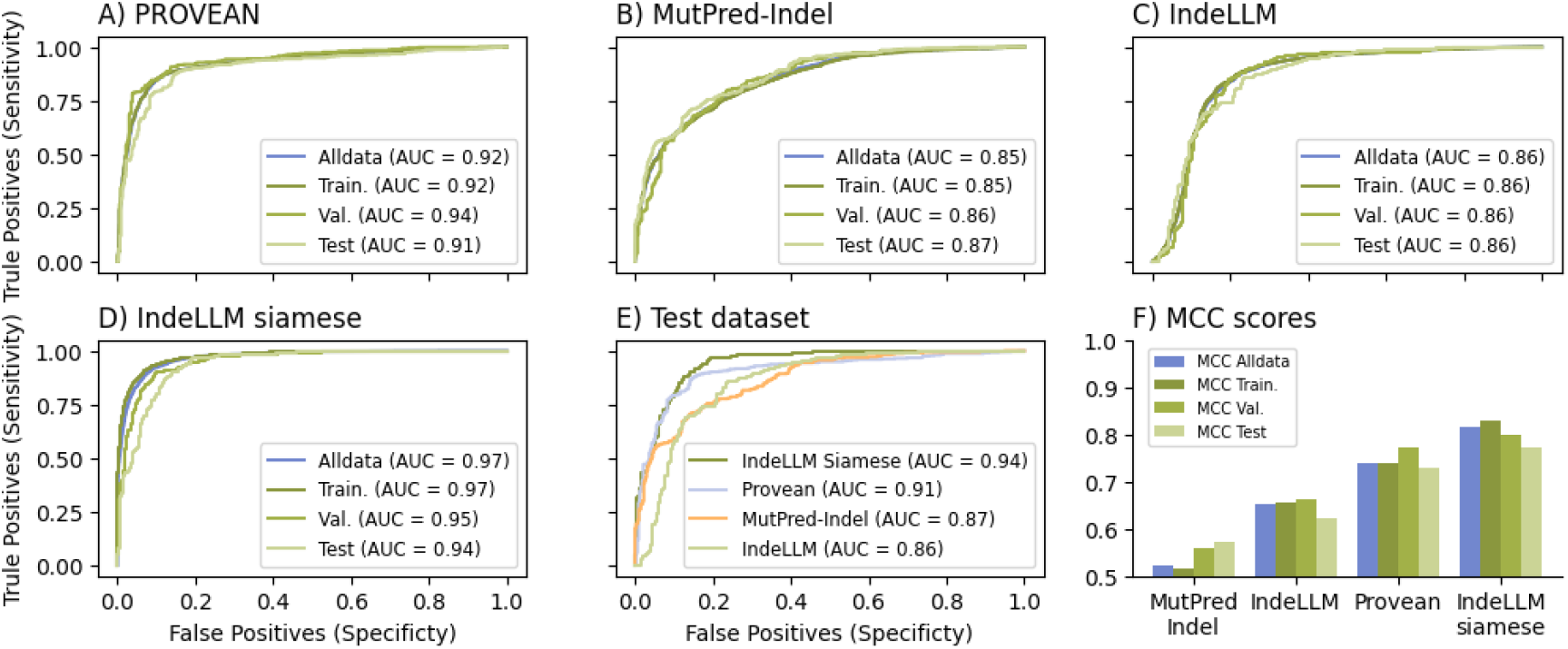
Evaluation of the performance of the best-performing pathogenicity prediction tools A) Provean and B) MutPred-indel, compared to C) IndeLLM and D) IndeLLM siamese on all collected indels (blue) and the different dataset splits (green) used for training (train.), validation (val.) and testing (test). In panel E, we compare their performance on the test dataset. In panel A-E, we plotted the ROC curve and calculated the AUC, while in panel F, we show the MCC scores for each method per dataset grouping and sorted by the MCC on the test data split.

The Siamese network outperforms the zero-shot inference by 24%. Examining the confusion matrix (Fig 4) reveals a high reduction in false negatives for insertions (19.51 % for IndeLLM, while 6.10% for IndeLLM siamese) (Fig 4B and 4D). For the IndeLLM score predictions, we observe that the false negative insertions have a high mean probability for the inserted amino acid, similar to the real benign insertions (Supplementary Materials, Section 5). Additionally, these false negative insertions tend to be longer than benign insertions. However, this effect cannot be observed using the IndeLLM Siamese model. This suggests that some insertions may not introduce damaging amino acids but may instead cause pathogenicity through increased stability or altered fit, which the IndeLLM scoring approach fails to detect. By incorporating sequence embeddings for the inserted or deleted amino acids in Siamese Model 4, we were able to account for these instances and accurately predict pathogenicity.

**Figure 4:**
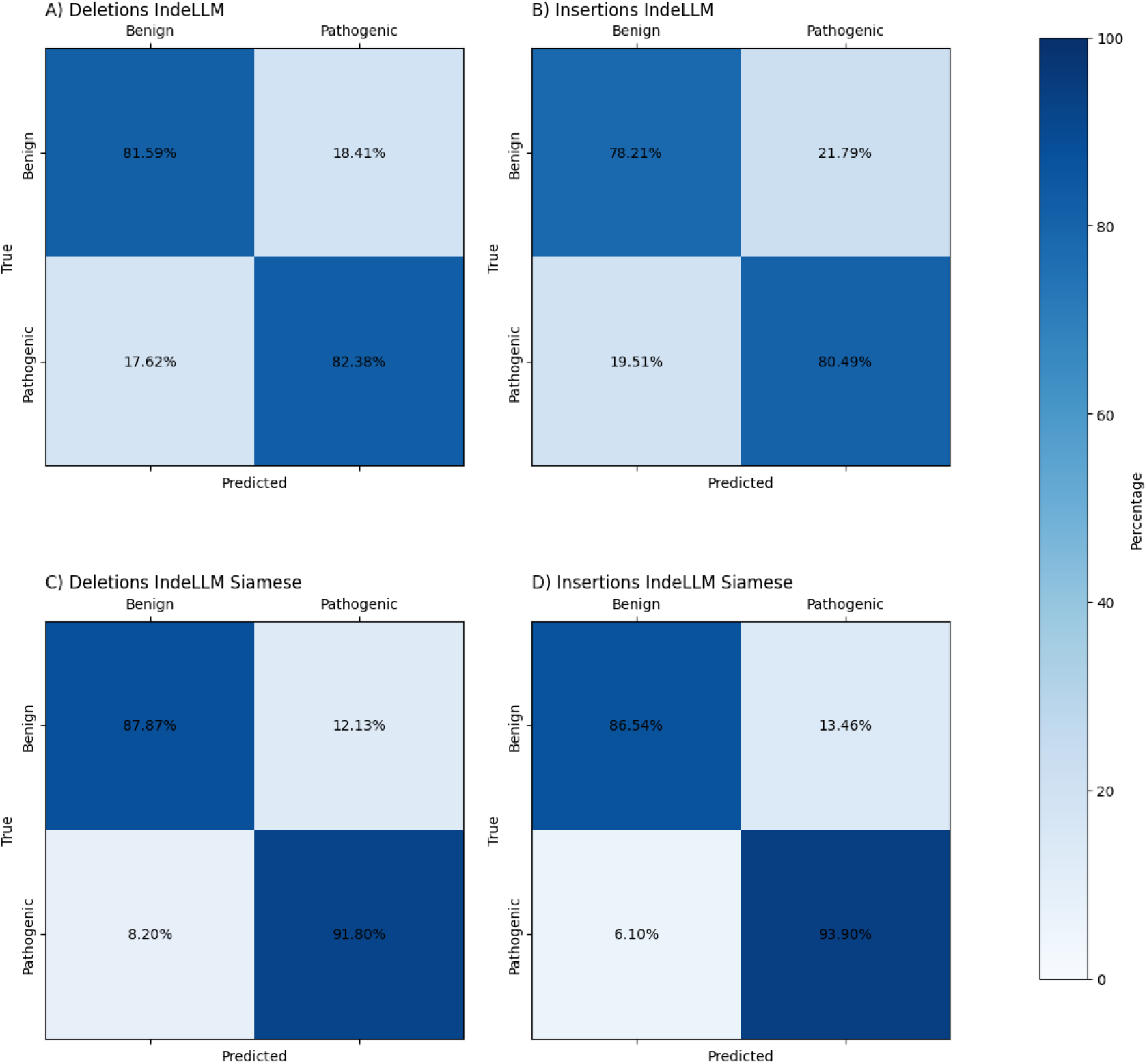
A) Confusion matrix for all deletions using IndeLLM scoring. B) Confusion matrix for all insertions using IndeLLM scoring. C) Confusion matrix for all deletions using IndeLLM siamese network, D) Confusion matrix for all insertions using IndeLLM siamese network. All confusion matrixes are coloured by the percentage of true positives for benign (left) and pathogenic (right) insertions or deletions.

### 4.4 Towards interpretable Indel predictions

Interpretability is essential for the usability of machine models. Understanding how these models predict changes in protein fitness allows users to validate results, ensuring that the predictions align with biological principles and are not artefacts or model biases.

The scoring method by PLMs per amino acid allows us to obtain information on the consequences of the indels for the protein environment. The model assigns a value to each amino acid in the sequence based on its predicted likelihood given the rest of the sequence, which indicates the evolutionary patterns. By subtracting the per amino acid wildtype probabilities from the per amino acid mutated probabilities (except the deleted or inserted amino acids) for the entire protein sequence, we obtain a difference in value per amino acid. The differences range from 1 to −1, where 1 can be interpreted as the indel introducing beneficial consequences for the amino acids, while −1 is detrimental. By plotting the differences in amino acid values, we can visualise the impact of the indel on the protein environment and, therefore, gain a better insight into why the indel is predicted as pathogenic or benign.

Our dataset reports four indels for a single protein, the Fibroblast growth factors receptor 1 (FGFR1). FGFR1 is a cell surface membrane receptor built up by three extracellular immunoglobulin-like domains (D1 – D3), a short hydrophobic transmembrane region, and a cytoplasmic tyrosine kinase domain [43]. Of the four indels, one is in the tyrosine kinase domain, and three are in the extracellular Ig-like domains (Table 2). The differences in amino acid probabilities illustrate which regions the indels impact (Fig 5A-5D). The two pathogenic deletions have a negative effect on the surrounding regions, while the two benign indels do not affect the protein. By colouring the 3D model by the difference, we provide an explanation for the prediction. Here, we show that deleting a Methionine on position 535 is predicted to destabilise the aC-helix (Fig 5E), a structural element essential during the activation of the enzyme [43]. We also found that deleting five amino acids in the first extracellular Ig-like domain (D1) in FGFR1. The long deletion occurs in a loop region, and IndeLLM predicts a complete destabilisation of the domain since we observe several interactions predicted to be altered (lost interactions in red and new interactions in blue Fig 5F). We tested our hypothesis by predicting the structure of the mutated domain using AlphaFold [44], which also predicts an unfolding of the domain (Supplementary Material, section 6)

**Figure 5:**
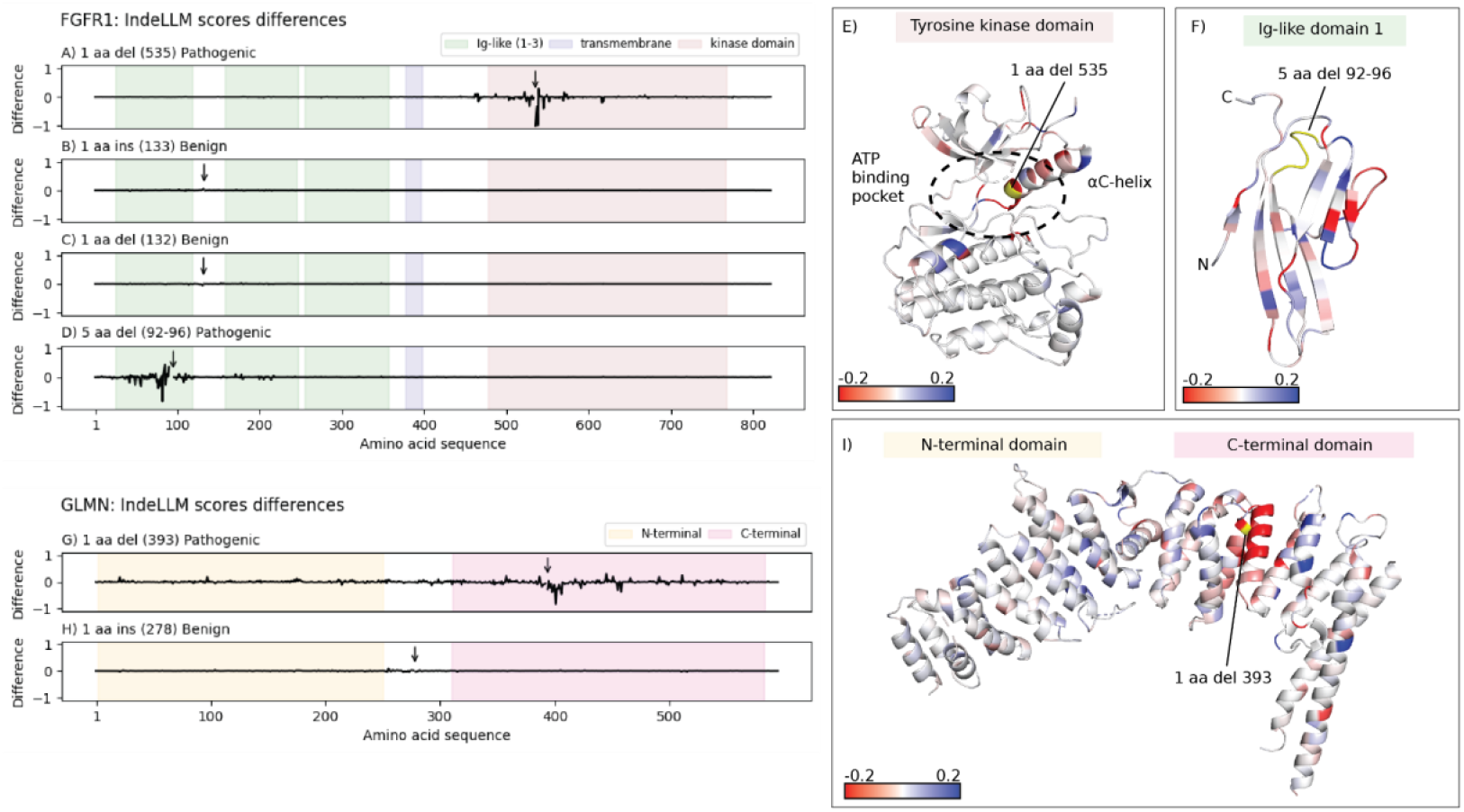
A-D) The difference between the wildtype and mutated sequence probabilities (y-axis) are plotted per amino acid in FGFR1 (x-axis). The coloured boxes on the plots represented the domains in FGFR1 (D1-D3 in green, transmembrane region in blue and the tyrosine kinase domain in red). In each plot, the indel position is labelled with an arrow and the indel type is noted in the title. E) PDB structure of the tyrosine kinase domain (PDB ID: 4UWY) coloured by the difference score (from red (<= −0.2) to blue (>= 0.2). The indels are labelled and coloured yellow. The impacted structural element (aC-helix) is labelled, as well as the ATP binding pocket (dotted circle). F) PDB structure of the Ig-like domain 1 (PDB ID: 2CR3) using the same colouring scheme as in panel E. The N and C-terminus are labelled. G-H) The plots are the same as in panel A-D, but for the GLMN protein sequence. The coloured boxes represented the N-terminal (yellow) and C-terminal (pink) domains. I) PDB structure of the GLMN protein (PDB ID: 4F52) is coloured as in panel E. The N- and C-terminal domains are labelled in coloured boxes above the structure.

**Table 2:** Example of indels reported for the FGFR1 protein (UniProt ID: P11362)

| Protein | Indel type | aa pos | FGFR1 domain | Wt sequence | Mut sequence | Indel size | ClinVar | IndeLLM |
| --- | --- | --- | --- | --- | --- | --- | --- | --- |
| FGFR1 | deletion | 535 | Tyrosine kinase domain | MK | K | 1 | Pathogenic/Likely pathogenic | Pathogenic |
| FGFR1 | insertion | 133 | Disordered | D | DD | 1 | Likely benign | Benign |
| FGFR1 | deletion | 132 | Disordered | DD | D | 1 | Likely benign | Benign |
| FGFR1 | deletion | 92 | Ig-like D1 | VPADSG | G | 5 | Likely pathogenic | Pathogenic |

Next we considered the effects of indels in Glomulin (GLMN), which is essential for normal vasculature development and is part of a large protein complex (Glomulin-RBX1-CUL1 Complex) [40]. GLMN (HEAT-like) adopts a repeated fold of a-helixes organised in N- and C-terminal domains. Our dataset reports two indels in the GLMN protein (Table 3). The pathogenic deletion is in the C-terminal domain, a domain that is essential for interactions with subunits in the complex [40]. The difference in probabilities between the wildtype and mutated sequence demonstrates that the deletion of Asparagine at position 393 destabilises large regions of the protein, particularly in the C-terminal domain (Fig 5G and 5I). In contrast, the benign insertion downstream of the N-terminal domain has minimal implications for the protein (Fig 5H).

**Table 3:**
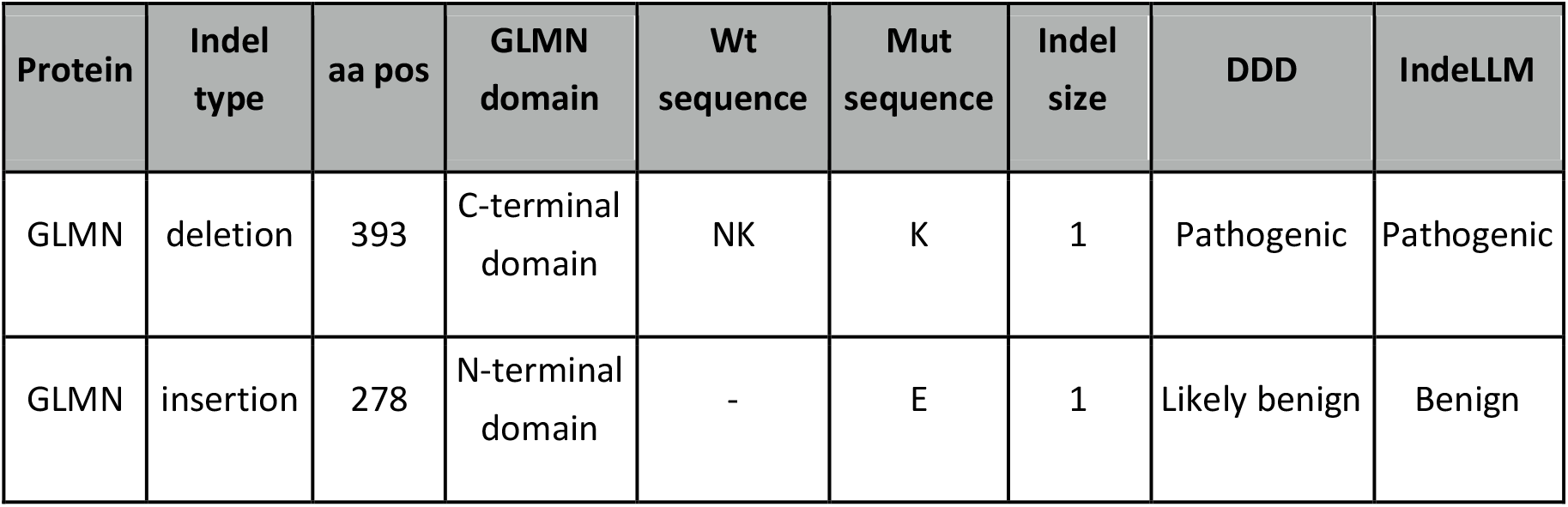
Example of indels reported for the GLMN protein (UniProtID: Q92990)

To show the easy use and interpretability of our scoring approach, we have designed a Colab available https://colab.research.google.com/drive/1CgwprttaNFR_KeJGyFzP0a0C9Y wc4P where any indel can be run independently of protein length, protein type and source organism, and the difference scoring plots (as shown in Fig 5) are automatically produced.

## 5. Discussion

Predicting the pathogenicity of insertions and deletions (indels) poses unique challenges due to their variable lengths and the diverse ways they can impact protein structure and function. Indels can result in different effects at the gene level, such as splicing changes, loss of function, frameshifts or in-frame mutations. Indels leading to large structural changes in the overall protein are generally damaging, but indels that preserve the open reading frame (in-frame indels) are especially challenging to characterise. In addition, the availability of labelled data for indels is far more limited than what is available for SNVs despite the relatively high abundance of indel mutations in the populations. This scarcity of high-quality data increases the risk of overfitting when training models, as they may learn patterns that are not generalizable to new cases.

Recent advancements in protein language models (PLMs), which have achieved significant success in protein structure prediction, functional annotation, and SNV pathogenicity assessment, constitute a promising foundation to tackle indel pathogenicity. These models are trained on massive protein sequence datasets, allowing them to capture complex sequence patterns and evolutionary relationships without the need for labelled data, making them potentially well-suited to interpret the intricate effects of indels, both as zero-shot methods or as features for downstream predictors. Efforts have been made to adapt these PLMs to the challenging task of indel pathogenicity prediction. However, the field of PLMs for indel prediction is still in its early stages, and there is room for further improvements. Additionally, up to this point, there are no techniques to aid in the interpretability of the results obtained from PLMs. Using PLMs as a zero-shot predictor utilising a scoring function is of high interest since it constitutes a straightforward approach with no risk of overfitting to known labelled data. Moreover, zero-shot approaches based on PLMs, since they do not rely on labelled data, can also be utilised for other organisms beyond humans, for which there may not be any information on pathogenicity.

The difference in length between the indel mutant and wildtype sequences makes for a non-straightforward scoring compared to SNV variants. A score for SNVs can be estimated by computing the log2 between the probabilities of the wildtype amino acid divided by the probability of the amino acid found in the mutant sequence. Brandes et al. developed a generalised scoring function for indels using PLMs based on calculating PLLs (Pseudo-log-likelihoods) of each of the sequences by adding the probability of each amino acid observed and then subtracting the PLL of the mutant and the wildtype sequence. One limitation of such an approach is that the PLLs used are not comparable due to the difference in length. The authors acknowledge the limitation in their original work and try to mitigate it by scaling the PLLs based on the difference in length. However, they observed no improvement compared to using the non-comparable PLLs directly and concluded that the PLLs obtained are robust enough to be used directly for predictions.

In this work, we have delved into the limitations and artefacts of using non-comparable PLLs. Due to the difference in length, we show that deletions are usually assigned higher pathogenicity scores than they should. At the same time, insertion received a lower one (mean Brandes score for deletions: −4.38 and insertions: 2.26). This difference in score comes from the extra amino acids in either the mutant or the wildtype sequence and the additive nature of this scoring function. We suggest a novel scoring approach that resolved this issue. We utilize the context-awareness nature of PLMs, which means that the effects of the insertions or deletions are indirectly inferred. We do this by calculating the PLLs using only the amino acids present in both the wildtype and mutant sequences, ignoring the inserted or deleted ones. This approach, which we named IndeLLM scoring, substantially improved the predictions, boosting PLMs predictive power by 67% (MCC score) to levels comparable to those methods trained with labelled data and manually curated features.

One appealing use of PLMs is to use their latent representation of the sequences as input features for downstream models. Various strategies have been proposed to integrate PLM embeddings across applications, including the prediction of insertions and deletions [19,23]. Ideally, such models should maintain simplicity to minimise the risk of overfitting when working with limited datasets. In this study, we explored the potential of a simple Siamese network architecture consisting of a single fully connected hidden layer with 8 nodes. Despite its simplicity, this architecture achieved state-of-the-art performance with no evidence of overfitting (IndeLLM siamese vs. Provean AUC: 0.94 vs. 0.91). To better understand the contributions of specific features, we conducted an ablation study, revealing that splitting the embedding of indel amino acids from the rest of the sequence embedding significantly improved performance. This underscores the value of incorporating biological insights into feature engineering which enhances model performance without increasing complexity. The proposed Siamese model, termed IndeLLM Siamese, outperformed previous approaches in classification accuracy (MCC 0.77) and demonstrated greater computational efficiency compared to previously suggested methods (Supplementary Material section 7). These findings indicate that the IndeLLM Siamese model can serve as a valuable tool for the community, both as a high-performing predictor and an example of effective feature selection in transfer learning.

A closer examination revealed that the improvements in the IndeLLM siamese, compared to the zero-shot IndeLLM scoring, come from the reduction of the insertions that were wrongly predicted as benign (false negatives). The confusion matrix for IndeLLM reveals a high false negative rate for insertions (19.51%), suggesting a systematic error in the predictions for insertions. A closer examination revealed that the majority of the false negative predictions of insertions had inserted amino acids with higher estimated probabilities than the average observed for pathogenic insertions (Supplement Material section 5). These finding suggests that such insertions could be causing a pathogenicity effect due to increase stability or altered fit, or perhaps gain-of-function [19]. IndeLLM score assesses the potential damage caused by the indel on the protein and uses that as a proxy for pathogenicity. This can explain why gain-of-function cases are not captured properly. We mitigated this by adding the information about the inserted or deleted amino acids in Siamese model 4 (Figure 2). The negative rate for insertions dropped drastically to 6.10%. By comparing the mean inserted probabilities of the false negative predicted insertions from IndeLLM and IndeLLM siamese model 4, we show a drop from 110 cases to 3 cases where the inserted amino acid had a higher probability than 0.8 (Supplement Material secion 5). This indicates that the IndeLLM siamese can capture gain-of-function cases using embeddings from the inserted amino acids.

Finally, we present a new approach to interpreting the models by examining individual amino acid probability changes. The obtained differences can then be mapped back to the structure to spot regions predicted to be affected by the indels. This allows experts to use this information, combined with their expertise on the protein in question, to examine more challenging cases of indels or spot other regions of interest that may be related to the indel at hand.

In summary, this work constitutes an in-depth analysis of the performance of PLMs of indels, highlighting current limitations to accurately predict pathogenic insertions caused by gain-of-function, as well as suggestions on improving PLM reliability. The improved scoring function allows for indel prediction performances on par with the best-performing methods without requiring any labelled data, minimising the risk of overfitting. Our approach is available to proteins from any organism, comes at a lower computational cost and faster assessment of variants compared to existing methods. We believe this information can aid in developing the indel predictor field. In line with that goal, much of our attention has been placed on making the method easy to use and accessible. The approach is freely available in Google Golab https://colab.research.google.com/drive/1CgwprttaNFR_KeJGyFzP0a0C9Y wc4P and the code and predictions used in this paper can be found in the following GitHub https://github.com/OriolGraCar/IndeLLM.

## 7. Supplementary

### Supplement File 1

- Table S1: MCC, F1 and AUC for scoring approaches
- Table S2: MCC, F1 and AUC for Siamese network models 1-4
- Table S3: MCC, F1 and AUC for IndeLLM, IndeLLM Siamese, Provean and MutPred-indel

### Supplement Material

- Section 1: Dataset curation and distribution of indel sizes
- Section 2: Comparing PLMs
- Section 3: Comparing Brandes and IndeLLM scores to nine prediction tools
- Section 4: Evaluating the level of overfitting
- Section 5: False negative insertions for IndeLLM and IndeLLM Siamese
- Section 6: AlphaFold prediction in mutated Ig-like domain
- Section 7: Computational efficiency

## 8. Data aviliablility

The source code for IndeLLM, all anlaysis, examples are available in our GitHub repository (https://github.com/OriolGraCar/IndeLLM). The IndeLLM Colab is aviliable at https://colab.research.google.com/drive/1CgwprttaNFR_KeJGyFzP0a0C9Y wc4P.

## 9. Acknowlegdment

The authors acknowledge The Research Council of Norway grant numbers 335244 and 350231 for funding toward running costs, travel grants, and conference support.

The authors declare no conflicts of interest.

