## Supplementary information for "Leveraging protein language models and scoring function for Indel characterisation and transfer learning"

### Supplementary Material

#### 1. Dataset curation

In this work, we publish a dataset of 7500 indels. These indels are combined from three studies (Cannon et al. [1], Fan et al. [2] and Brandes et al. [3]). IndeLLM requires protein sequences as input. Using gene and transcriptions IDs, we collected the protein sequences for the indels where this information was not included. The curation of all indels (ensuring correct protein sequence, indel annotation, excluding any frameshift or delins (which are deletions followed by insertions resulting in identical length of wildtype and mutated sequence)) resulted in the exclusion of hundreds of indels. However, the final dataset of 7500 indels is thoroughly curated and represents, to our knowledge, the largest dataset of in-frame indels with pathogenicity classifications. The dataset includes 2409 insertions and 5091 deletions. Of these, 2878 are classified as likely pathogenic or pathogenic, while 4622 are classified as likely benign or benign. The indels range from 1 to 223 in length and represent a distribution of short indels (1-2 amino acid, n = 3882) and longer indels (>= 3 amino acids, n = 3618). See Figure S1 for distribution.

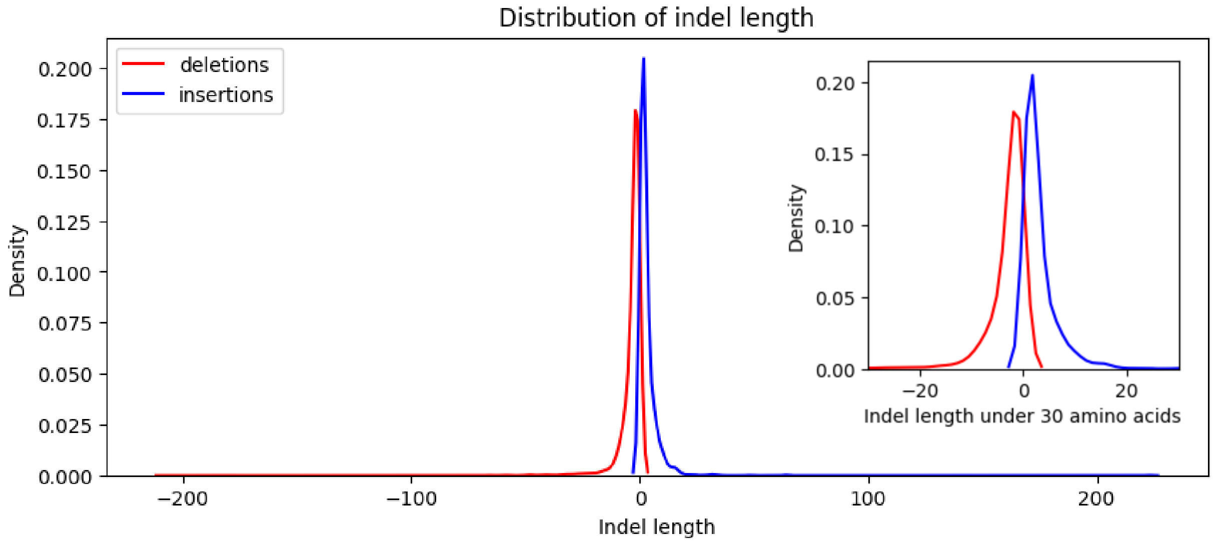

*Figure S1: The distribution of indels by length (deletions in red and insertions in blue).*

#### 2. Comparing 7 PLMs

We have studied the performance and transferability of 7 PLMs, ESM1v [4], ESM1b [5], ProtTrans [6], ESM2 at three different sizes (150M, 650M and 3B parameters) [7] and ESM3 (1.4B parameters) [8]. All PLMs were run using the HuggingFace transformers library and the available pre-trained parameters from Huggingface, except ESM3, which was run using the code available on the official GitHub repository of the Model (https://github.com/evolutionaryscale/esm). ESM3 is a multimodal generative language model that can be inputted with information for different tracks. This work's goal was to examine the performance of language models using protein sequence alone, so only the protein track was used to assess the performance of ESM3.

Using the 7500 indels dataset and our IndeLLM score, we found that three PLMs performed the best (ESM1b, ESM2-650M, and ESM2-3B), with similar accuracies: MCC of 0.68 vs. 0.65 vs. 0.64, respectively (Figure S2 and Table S1). Due to the flexibility of rotatory positional embedding, ESM2-650M is the best candidate for encoding longer sequences without extra preprocessing steps.

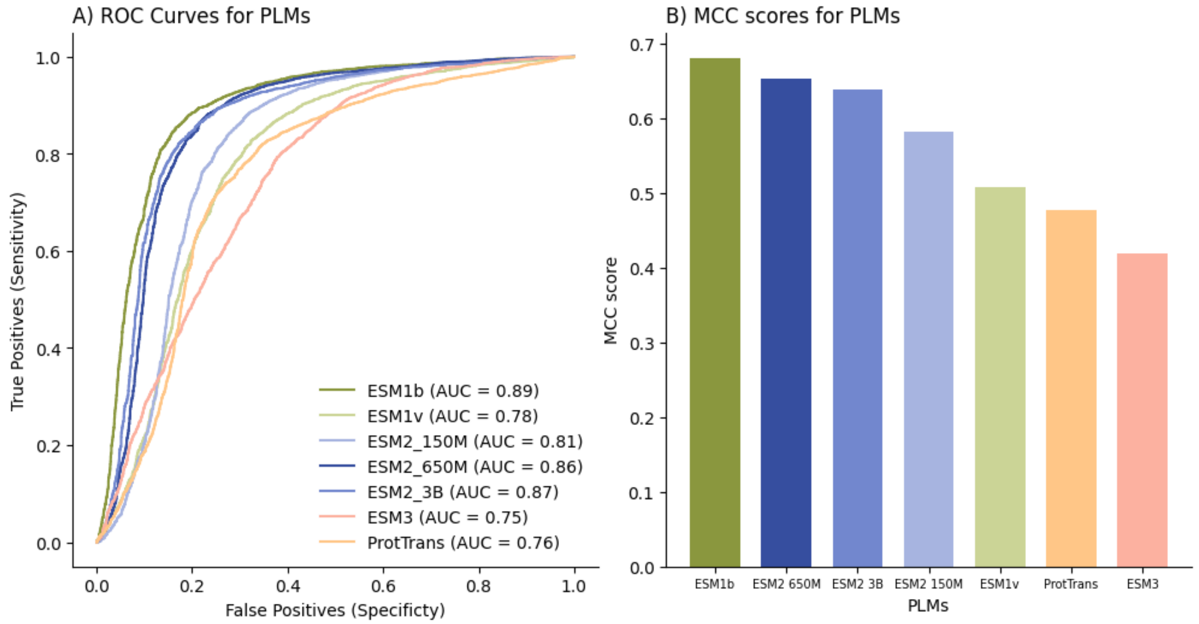

*Figure S2: Evaluation of different PLMs using the local scoring approach. The different PLMs are ESM1 (b and v, green), ESM2 (150M, 650M and 3B parameters, blue), ESM3 (orange) and ProtTrans (red). We compared the scoring methods by A) plotting the ROC curve and calculating the AUC and B) calculating the MCC scores.*

Full dataset (n = 7500)

| PLMs* | MCC | F1 | AUC |
|---|---|---|---|
| ESM1b | 0.68 | 0.87 | 0.89 |
| ESM2 (650M paramteres) | 0.65 | 0.87 | 0.86 |
| ESM2 (3B paramteres) | 0.64 | 0.85 | 0.87 |
| ESM2 (150M paramteres) | 0.58 | 0.85 | 0.81 |
| ESM1v | 0.51 | 0.82 | 0.78 |
| ProtTrans | 0.48 | 0.80 | 0.76 |
| ESM3 | 0.42 | 0.78 | 0.75 |

*Table S1: The calculated MCC, F1 and AUC scores for the 7 PLMs (*All results use indeLLM score).*

Models with additional parameters, such as ESM2-3B (MCC score 0.64), did not perform better than the smaller models, suggesting that these smaller checkpoints may be more suitable to represent indels. The trend of bigger models not always correlating with better performance has also been previously observed in the pure Large Language Models (LLMs) [9]. ESM3 performed worst among the tested ESM models (MCC 0.42). However, it is important to note that we only used the sequence track in our analysis, which leaves the question of how ESM3 would perform if extra features were provided in the other tracks.

#### 3. Comparing Brandes and IndeLLM scores to nine prediction tools

Cannon et al. 2023 [1] include pathogenicity prediction from nine tools: CADD, CAPICE, FATHMM-indel, MutPred-Indel, MutationTaster2021, PROVEAN, SIFT-indel, VEST-indel, and VVP. We used the Cannon et al. subset in the full dataset, which holds 3478 indels from gnomAD (n = 647), ClinVar (n = 2577), and DDD (n = 254), where 1115 are insertions and 2363 are deletions. Of these, 1518 are classified as likely pathogenic or pathogenic, while 1960 are classified as likely benign or benign. We compared performance using the Brandes and IndeLLM score to the nine indels predictors reported (Figure S3 and Table S2). Brandes score underperformed, ranking last among the tested methods (MCC score: 0.38), while using the IndeLLM score resulted in a performance with similar accuracy to the best-performing predictors (MCC scores: IndeLLM = 0.69, MutationTaster2021 = 0.93, PROVEAN = 0.77 and Vest-indel = 0.74).

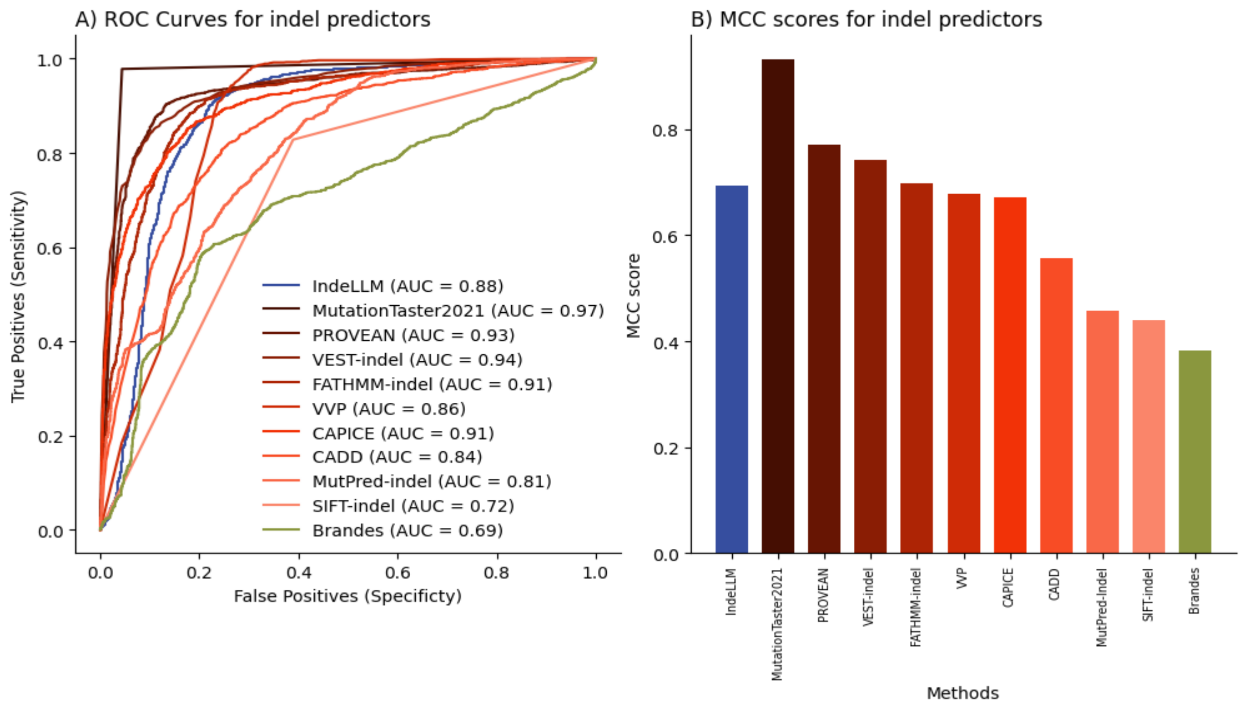

*Figure S3: Evaluation of different prediction tools (red) to IndeLLM score (blue) and Brandes Score (green). We compared the scoring methods by A) plotting the ROC curve and calculating the AUC and B) calculating the MCC scores.*

Cannon et al. dataset (n = 3478)

| **Published indel predictors** | **MCC** | **F1** | **AUC** |
|---|---|---|---|
| MutationTaster2021 | 0.93 | 0.97 | 0.97 |
| PROVEAN | 0.77 | 0.90 | 0.93 |
| VEST-indel | 0.74 | 0.89 | 0.94 |
| FATHMM-indel | 0.70 | 0.86 | 0.91 |
| VVP | 0.68 | 0.83 | 0.86 |
| CAPICE | 0.67 | 0.85 | 0.91 |
| CADD | 0.56 | 0.79 | 0.84 |
| MutPred-indel | 0.46 | 0.72 | 0.81 |
| SIFT-indel | 0.44 | 0.70 | 0.72 |
| **Scoring method*** | | | |
| IndeLLM | 0.69 | 0.87 | 0.88 |
| Brandes | 0.38 | 0.67 | 0.69 |

*Table S2: The calculated MCC, F1 and AUC scores for the prediction tools and IndeLLM and Brandes scores (*Scoring methods results reported are calculated using the probabilities from ESM2 (650M parameters)).*

#### 4. Evaluating the level of overfitting

Cannon et al. suggested using the DDD indels (n=254) [1] as a good candidate to examine overfitting on different methods with labeled data since the dataset was made available after those methods were published. We expanded the original analysis of overfitting to all the reported methods (Figure S4 and Table S3). Several methods displayed degrees of overfitting. Among them was the best-performing method, MutationTaster2021, having its performance cut from MCC 0.95 to 0.51 (Fig 4E and 4L). No overfitting was observed for PROVEAN and MutPred-Indel (MCC dropped 0.11 and 0.10, respectively, Fig 4D, 4F, and 4L). As expected, the performance of IndeLLM and Brandes showcased no overfitting since no indel sequences were used at any point to train or readjust any of the original weights (Fig 4J, 4K and 4L).

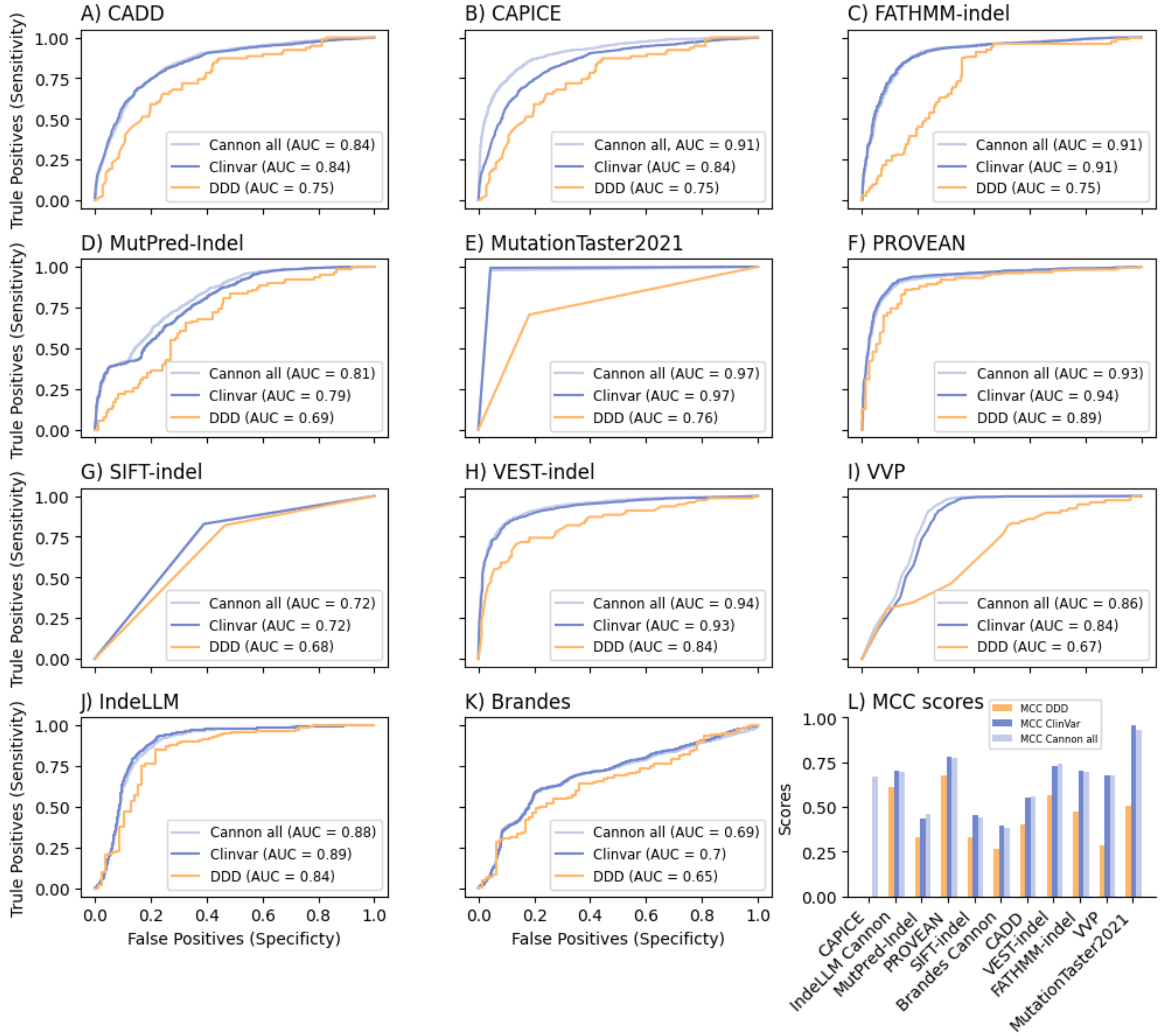

*Figure S4: Evaluation of the level of overfitting of pathogenicity prediction tools. Panel A-I shows the AUC for nine prediction tools. Panel J shows the AUC for IndeLLM scoring. Panel K shows the AUC for Brandes scoring. We calculate AUC using all Cannon et al. indels (light blue), the Cannon et al. indels collected from ClinVar (dark blue) and the Cannon et al. indels collected from DDD*

*(orange). In panel L, we show the calculated MCC scores for each method per dataset split and sorted by the difference between MCC ClinVar and MCC DDD. CAPICE has no calculations of MCC for the ClinVar and DDD data splits since the tool predicts all indels as either pathogenic or benign, making the binary prediction one-dimensional.*

Cannon et al. Dataset split by indel origin, ClinVar (n = 2577) and DDD (n = 254)

| | **ClinVar** | | | **DDD** | | | **DDD - ClinVar** |
|---|---|---|---|---|---|---|---|
| **Published indel predictors** | **MCC** | **F1** | **AUC** | **MCC** | **F1** | **AUC** | **MCC diff** |
| MutPred-Indel | 0.43 | 0.65 | 0.79 | 0.33 | 0.65 | 0.69 | 0.10 |
| PROVEAN | 0.78 | 0.88 | 0.94 | 0.68 | 0.89 | 0.89 | 0.11 |
| SIFT-indel | 0.45 | 0.67 | 0.72 | 0.33 | 0.66 | 0.68 | 0.12 |
| CADD | 0.55 | 0.75 | 0.84 | 0.40 | 0.69 | 0.75 | 0.15 |
| VEST-indel | 0.73 | 0.85 | 0.93 | 0.57 | 0.87 | 0.84 | 0.16 |
| FATHMM-indel | 0.70 | 0.83 | 0.91 | 0.48 | 0.75 | 0.75 | 0.22 |
| VVP | 0.67 | 0.79 | 0.84 | 0.28 | 0.61 | 0.67 | 0.39 |
| MutationTaster2021 | 0.95 | 0.97 | 0.97 | 0.51 | 0.84 | 0.76 | 0.45 |
| CAPICE | 0** | 0.61 | 0.84 | 0** | 0.82 | 0.75 | 0.67 |
| **Scoring method*** | **MCC** | **F1** | **AUC** | **MCC** | **F1** | **AUC** | **MCC diff** |
| IndeLLM | 0.70 | 0.84 | 0.89 | 0.61 | 0.87 | 0.84 | 0.09 |
| Brandes | 0.40 | 0.64 | 0.70 | 0.27 | 0.62 | 0.65 | 0.09 |

*Table S3: The calculated MCC, F1 and AUC scores for the prediction tools and IndeLLM and Brandes scores for the ClinVar and DDD data splits. The final column calculates the difference in MCC (*Scores are calculated using the probabilities from ESM2 (650M parameters), **MCC not calculated due to CAPICE binary predictions being one-dimensional).*

#### 5. False negative insertions for IndeLLM and IndeLLM Siamese

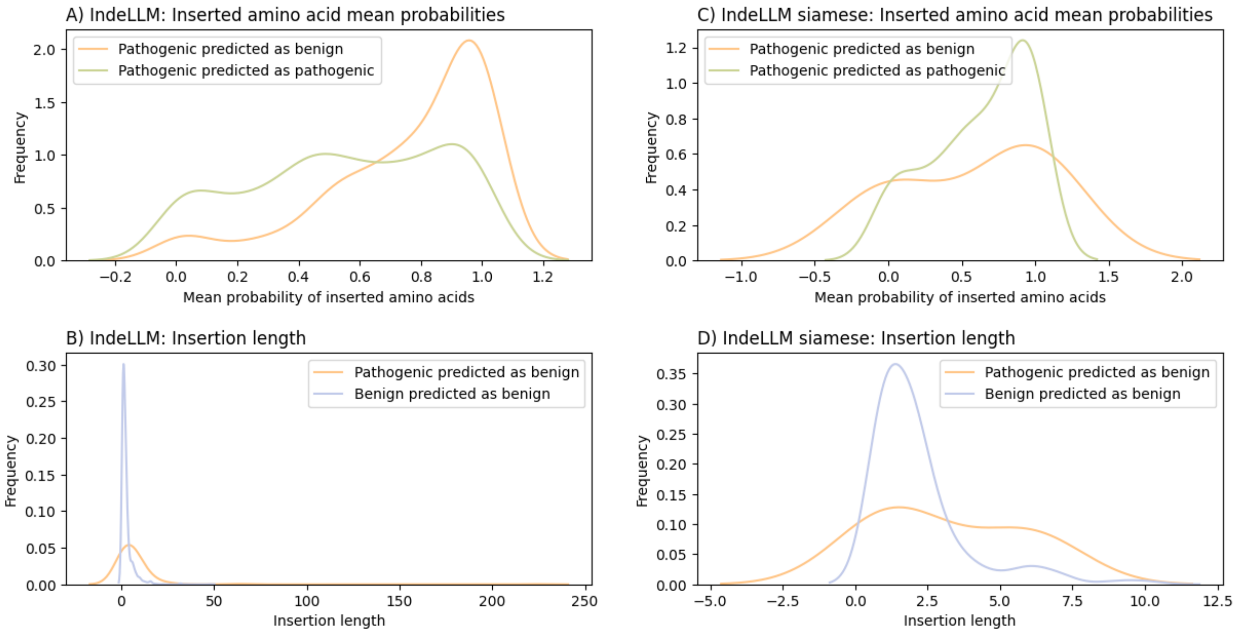

*Figure S5: Panels A and B show IndeLLM score calculations, while panels C and D are from IndeLLM Siamese. Panels A and C show the mean value of inserted amino acid, which was calculated for true positives (green) and false negatives (orange). In contrast, panels B and D show the length of inserted amino acids, which were calculated for false negatives (orange) and true positives (blue).*

#### 6. AlphaFold prediction

To test our hypothesis that the 5 amino acid deletion in the Ig-like D1 domain of FGFR1 results in destabilisation of the domain, we ran the mutated sequence of the Ig-like domain (Uniprot ID: P11362 residue 37 to 125) on AlphaFold Collab [10]. All predictions from AlphaFold predict loss of a β-strand (Figure S6). The loss of this secondary structure element is most likely destabilising for the domain structure.

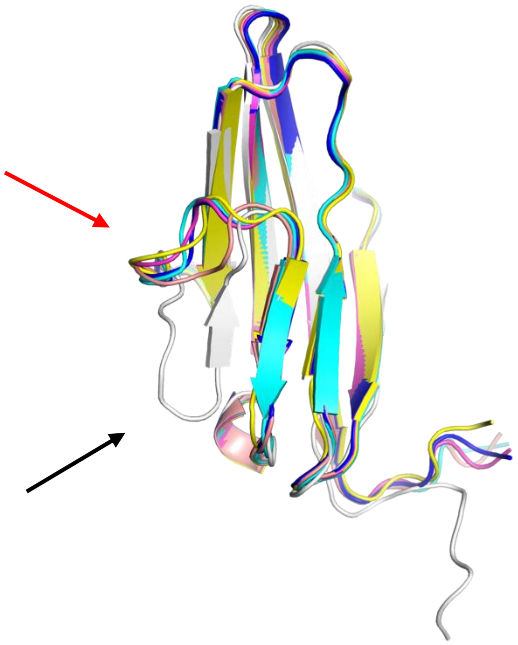

*Figure S6: The wildtype Ig-like domain (PDB ID: 2CR3) superimposed on the five AlphaFold models (yellow, cyan, magenta, pink and blue). The black arrow highlights the lost β-strand in all predicted models (wildtype in white). The red arrow points to the truncated loop in the AlphaFold predictions.*

#### 7. Computational efficiency

Provean is no longer supported (https://www.jcvi.org/research/provean), and the download is no longer compatible with today's systems. At the time of this analysis, the web server was accessible but allowed 100 indels runs per day, and processing of each indel takes 10-15 minutes if the sequences are not already downloaded in the server [11].

We downloaded the MutPred-Indel standalone to run the indels. We run all indels in batches of 100. The mean processing time per batch run was 4 hours and 35 minutes (2.75 minutes per job). However, some of this processing time is due to startup of the MATLAB dependency (http://mutpred2.mutdb.org)[12].

Non-multiple-sequence alignments (MSAs) Transformer-based approaches, as described in this work, are normally significantly faster than traditional methods that rely on multiple-sequence MSAs or data integration from multiple sources. One key advantage of transformers is their ability to process entire sequences in parallel using self-attention mechanisms rather than requiring iterative or sequential computations. This parallelisation allows transformer models to analyse protein sequences in batch, taking advantage of modern GPU and TPU architectures for high-throughput processing. A complete comparison of the performances of the two methods is out of the scope of this manuscript. However, using an RTX4000, IndeLLM was several orders of magnitude faster than the other methods that could be run locally.
